# An integrative analysis of GWAS and intermediate molecular trait data reveals common molecular mechanisms supporting genetic similarity between seemingly unrelated complex traits

**DOI:** 10.1101/601229

**Authors:** Jialiang Gu, Chris Fuller, Jiashun Zheng, Hao Li

## Abstract

The rapid accumulation of Genome Wide Association Studies (GWAS) and association studies of intermediate molecular traits provides new opportunities for comparative analysis of the genetic basis of complex human phenotypes. Using a newly developed statistical framework called Sherlock-II that integrates GWAS with eQTL (expression Quantitative Trait Loci) and metabolite-QTL data, we systematically analyzed 445 GWAS datasets, and identified 1371 significant gene-phenotype associations and 308 metabolites-phenotype associations (passing a Q value cutoff of 1/3). This integrative analysis allows us to translate SNP-phenotype associations into functionally informative gene-phenotype association profiles. Genetic similarity analyses based on these profiles clustered phenotypes into sub-trees that reveal both expected and unexpected relationships. We employed a statistical approach to delineate sets of functionally related genes that contribute to the similarity between their association profiles. This approach suggested common molecular mechanisms that connect the phenotypes in a subtree. For example, we found that fasting insulin, fasting glucose, breast cancer, prostate cancer, and lung cancer clustered into a subtree, and identified cyclic AMP/GMP signaling that connects breast cancer and insulin, NAPDH oxidase/ROS generation that connects the three cancers, and apoptosis that connects all five phenotypes. Our approach can be used to assess genetic similarity and suggest mechanistic connections between phenotypes. It has the potential to improve the diagnosis and treatment of a disease by mapping mechanistic insights from one phenotype onto others based on common molecular underpinnings.

## Introduction

Genome Wide Association Studies (GWAS) seek to identify the genetic basis of complex phenotypes by associating genetic variants (typically characterized by Single Nucleotide Polymorphisms, or SNPs) to particular traits across different individuals. These studies have been applied to a wide range of phenotypes, including many complex diseases, and have led to the identification of a large number of informative SNPs [1, 2]. GWAS findings have revealed significant insights into disease mechanisms and informed strategies for improved intervention and prevention. GWASs have also been successfully implemented to define the relative influence of genotype and environment on phenotype prevalence, assisting in risk prediction[1].

Despite these successes, traditional GWAS analyses based on the association of individual SNPs have only limited power. Many potentially informative SNPs fall below the genome-wide significance threshold, which is set high due to the large number of loci typically tested. Also, for associations that do exceed the genome-wide significance threshold, the functional link between an individual SNP and the phenotype is often poorly understood. To circumvent these problems, integrative analysis methods have emerged that combine GWAS with other quantitative molecular data. These methods attempt to bridge the knowledge gap between GWAS SNPs and phenotypes by uncovering the genes and other intermediate molecular traits (IMTs) through which genetic polymorphisms generate phenotypic differences [3–7]. These approaches utilize the collective power of SNPs and generate mechanistic hypotheses that can be tested and can guide future therapeutic interventions.

As GWAS have been applied to additional human traits, it has become increasingly apparent that seemingly unrelated complex traits may share common genetic origins[8–11]. Phenotypic connections (often in the form of co-occurrence) have long been observed based on epidemiological studies [12]. Recent comparative GWAS analyses have lent genetic support to these earlier observations by searching for pleiotropic SNPs implicated across multiple phenotypes[13–15]. However, genetic similarity between different phenotypes can be obscured at the SNP level. Phenotypes sharing a set of IMTs may not necessarily associate with the same SNPs, since multiple SNPs can converge on the same IMT, and only a small subset of these SNPs will be identified in a particular GWAS. Furthermore, it is quite difficult to infer common mechanisms based on the shared SNPs, as they often fall into non-coding regions with no obvious functional implication.

We developed an integrative analysis framework in which GWAS-IMT analysis is combined with comparative analysis of multiple phenotypes (Figure 1). Such analysis not only enables us to translate SNP-phenotype association to IMT-phenotype association (which yields improved statistical power and mechanistic insight), but also to generate hypotheses regarding common biological processes underlying different (and sometimes seemingly unrelated) phenotypes based on the similarity of their IMT association profiles.

**Figure 1.**
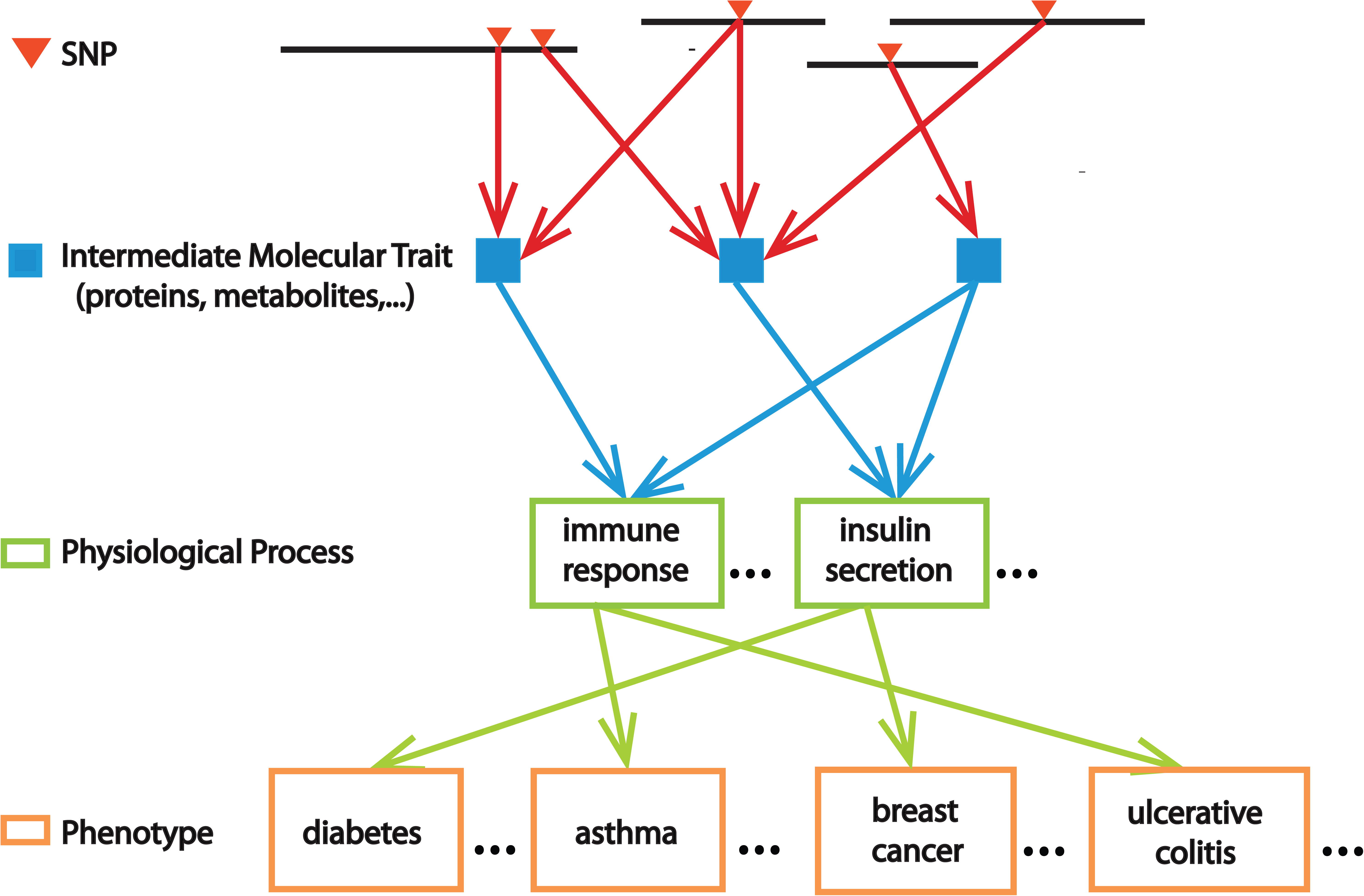
A schematic of the integrative analysis framework. The underlying model is that SNPs (red triangles) influence phenotypes (orange boxes) by influencing intermediate molecular traits (IMTs, blue squares) which in turn contribute to various physiological processes (green boxes), and each phenotype is controlled by a combination of physiological processes. In the first step of the analysis, data from GWASs and association studies of IMTs (such as eQTL and metabolite-QTL) are integrated to infer IMT-phenotype associations. A comparative analysis of the IMT association profiles for different phenotypes then identifies genetic similarity between phenotypes, supported by shared IMTs. These IMTs may point to a set of common physiological processes (blue arrows pointing to the same green boxes), suggesting shared physiological processes underlying different phenotypes (green arrows initiating from the same green box).

The first step in this integrative analysis is to associate the IMTs with phenotypes using all the SNPs present in GWAS results. We developed a new computational algorithm called Sherlock-II to integrate GWAS data with eQTL and metabolite-QTL data, translating SNP-phenotype associations into gene/metabolite-phenotype associations. This produces a gene-phenotype association matrix and a metabolite-phenotype association matrix and defines a gene association profile for each phenotype (one column of the gene-phenotype association matrix).

We then analyze the relation between phenotypes and infer possible common underlying physiological processes (PP) by two different approaches. In the first approach, we group the phenotypes based on the similarity between their gene association profiles and then identify groups of functionally related genes (such as those defined by gene ontology (GO) terms or pathways) that contribute to the similarity. This generate hypotheses on PPs (green boxes in Figure 1) that connect multiple phenotypes (orange boxes in Figure 1).

In the second approach, we apply two-way bi-clustering to the gene-phenotype association matrix to identified sub-clusters in which a group of genes are associated with a subset of phenotypes. We hypothesize that such a group of genes participate in the same PPs that influence the subset of phenotypes. We conducted literature searches on each group of genes to identify similar mechanisms through which they may affect the phenotypes, such as common pathways. This often leads to important clues into the PPs involved.

Our analysis predicts new connections between PPs and phenotypes (Figure 1) supported by multiple genes and thus generates etiological hypotheses with genetic support. Identifying different diseases associated with common PPs also provide an opportunity to transfer existing intervention from one disease to another.

## Results

### Sherlock-II: a new computational algorithm to infer IMT-phenotype association by integrating GWASs and association studies of IMT

To investigate the underlying biology of multiple phenotypes simultaneously, we need a high throughput tool to combine GWAS data with molecular trait association data in a statistically robust manner. To meet this need, we built a computational tool called Sherlock-II (a new generation of our previous published tool Sherlock[5]) to infer IMT-phenotype associations from GWAS and functional datasets. Similar to Sherlock, Sherlock-II detects gene-phenotype association by comparing the GWAS SNP with the eQTL SNP of a gene, with the assumption that if the gene is causal to the phenotype, SNPs that influence the expression of the gene should also influence the phenotype, thus the eQTL profile of the gene should have significant overlap with the GWAS profile of the phenotype. Sherlock-II uses a new statistical approach to directly calculate the statistical significance of the overlap, leading to a more efficient algorithm that is robust against inflation of the input GWAS data. By deriving the background distribution empirically from the list of all p-values for GWAS SNPs, it automatically accounts for inflation that is typically present in GWAS results (see Method). In addition, the statistical framework used by Sherlock-II is also directly applicable to metabolite-QTL data [16], allowing us to infer associations between metabolite levels and phenotypes.

### Global analysis using Sherlock-II identified many gene/metabolite-phenotype associations

Using Sherlock-II, we have systematically analyzed the published GWAS data, using a set of eQTL and metabolite-QTL data we curated. The analyses include 445 GWAS datasets covering 92 phenotypes, 8 individual eQTL covering 6 unique tissues[17–24], GTEx (version 7) data[25], and 1 metabolite-QTL dataset [14]. To include the possibility that the expression of a gene can influence the phenotype through multiple tissues, we combined eQTL data across different tissues: when a SNP appears in multiple eQTL datasets, we assign the most significant p-value to it in order to capture the best eQTL signal across tissues. This merged version includes the eQTL profiles of 21,892 genes. For each GWAS dataset, Sherlock-II calculates a p-value for every gene measuring the statistical significance of the overlap between the merged eQTL profile of the gene and the GWAS profile. Thus Sherlock-II translates the original SNP-phenotype association to gene-phenotype association. Similarly, by integrating metabolite-QTL data with GWAS data, Sherlock-II translates SNP-phenotype association to metabolites-phenotype association. Overall, we have identified 2145 gene-phenotype associations (with Q-value <1/3) covering 1371 genes and 74 phenotypes, and 482 metabolite-phenotype associations (Q<1/3) covering 308 metabolites and 53 phenotypes. In many cases, although only weak individual associations are present in the GWAS data, Sherlock-II detects strong gene/metabolite-phenotype association signals as many GWAS SNPs are considering in aggregate during the transformation. By combining across multiple SNPs with weak associations, information present in GWAS associations at levels below traditional gnome-wide significance thresholds can be extracted. Our global analysis uncovers many gene/metabolite-phenotype associations that shed light on the genetic determinants of complex human traits, including disease risks.

As an example, we have identified several genes significantly associated with rheumatoid arthritis (RA). Three out of five most significant RA associated genes are from human leukocyte antigen family (HLA-DRB5, HLA-DPA1, HLA-DPB2), consistent with the established immune origin of RA [26]. The third most significant gene ITPR3(inositol 1,4,5-Trisphosphate Receptor Type 3) mediates the release of intracellular calcium, and has been identified as RA associated gene through a pathway based analysis [27]. The involvement of intracellular calcium signaling in the RA pathology is supported by single cell imaging showing abnormal intracellular calcium signaling in RA patients[26]. We also identified CACNB1(Calcium Voltage-Gated Channel Auxiliary Subunit Beta 1), further supporting the role of abnormal calcium signaling in RA. In addition, two other genes involved in regulating apoptosis (BAK1 and GSDMB) are significantly associated with RA, and cytokine induced cell apoptosis were observed in RA synovium[28].

Application of Sherlock-II to integrate GWAS with metabolite-QTL data leads to the identification of 482 significant metabolites-phenotype associations. For example, we have replicated the association between methionine-sulfone and chronic kidney disease found in a previous study[16] with more supporting SNPs. Interestingly, metabolite-phenotype associations seem to be enriched for neuropsychiatric/neurodegenerative diseases (Fisher’s exact test p-value= 0.02215). This enrichment becomes more obvious when more stringent Q-value threshold is applied (Table S1). For examples, we have identified a strong association between gamma-glutamylglutamate and schizophrenia (p-value=3.97E-07, Q-value=2.46E-04); this metabolite was observed to decrease in schizophrenic patients and is one of five best discriminators between the schizophrenic patients and the controls[29]. We have identified palmitoyl-dihydrosphingomyelin as the most significantly associated metabolite with Alzheimer’s disease (p-value=7.38E-05, Q-value=4.45E-02). Palmitoyl-dihydrosphingomyelin is a particular type of sphingomyelin and it has been reported that a significant increase of sphingomyelin levels have been observed in the cerebrospinal fluid from individuals with prodromal Alzheimer’s Phenotype compared to cognitively normal controls [30]. For Autism, the most significantly associated metabolite is homocitrulline, and decreased homocitrulline concentration has been detected in Autistic patients[31]. In all the cases except methionine sulfone, the metabolite was implicated by multiple weak metabolite-QTL SNPs aligned to moderate GWAS SNPs, thus cannot be identified by genome-wide significant metabolite-QTL SNPs and GWAS SNPs alone (Figure S1).

The above results suggest that the significant gene/metabolite associations with other phenotypes identified by Sherlock-II are also likely to be biologically meaningful and detailed analysis can lead to new insight into the disease etiology. To facilitate the analysis by the community we have provided a full list of the gene/metabolite-phenotype associations in Table S2.

### Global analysis of the genetic similarity between phenotypes

The global gene-phenotype association matrix not only allows us to identify genes associated with a specific phenotype but also enables the analysis of the genetic similarity between different phenotypes at the gene level. Previous analyses of genetic similarity at the SNP level revealed certain overlapping SNPs between statistically associated phenotypes, lending genetic support to the observed co-occurence [32–34]. Genetic similarity at the gene level may yield stronger signal as multiple SNPs may converge at the same gene (e.g., influencing the expression of the same gene through different eQTL SNPs (eSNPs), thus one gene may be implicated in multiple phenotypes that do not necessarily share SNPs in the original GWAS data. Indeed, we have observed many instances where a gene’s eQTL profile aligned to the GWAS profiles of two different phenotypes through different SNPs (see Figure S2 for an example). More importantly, genetic similarity analysis at the gene level also allows us to delineate the common cell/molecular mechanisms underlying different phenotypes, which we describe in more detail in the next section.

We first seek to identify the global similarity between two different phenotypes by comparing the gene-phenotype association vector (−log10(p-value) for each gene, with a dimension of the number of genes = 21892) for the two phenotypes. Intuitively, similar gene p-value vectors imply that the phenotypes respond similarly to the expression changes of individual genes. We applied hierarchical clustering to cluster all the phenotypes, using Euclidean distance between gene-phenotype vectors. The result of this global clustering analysis is a tree in which phenotypes with similar p-value vector are placed next to each other in a subtree (Figure 2). We find similar/related phenotypes were clustered together as expected. However, more interestingly we find phenotypes that are not obviously related are also clustered together, suggesting that they share similar genetic mechanisms.

**Figure 2.**
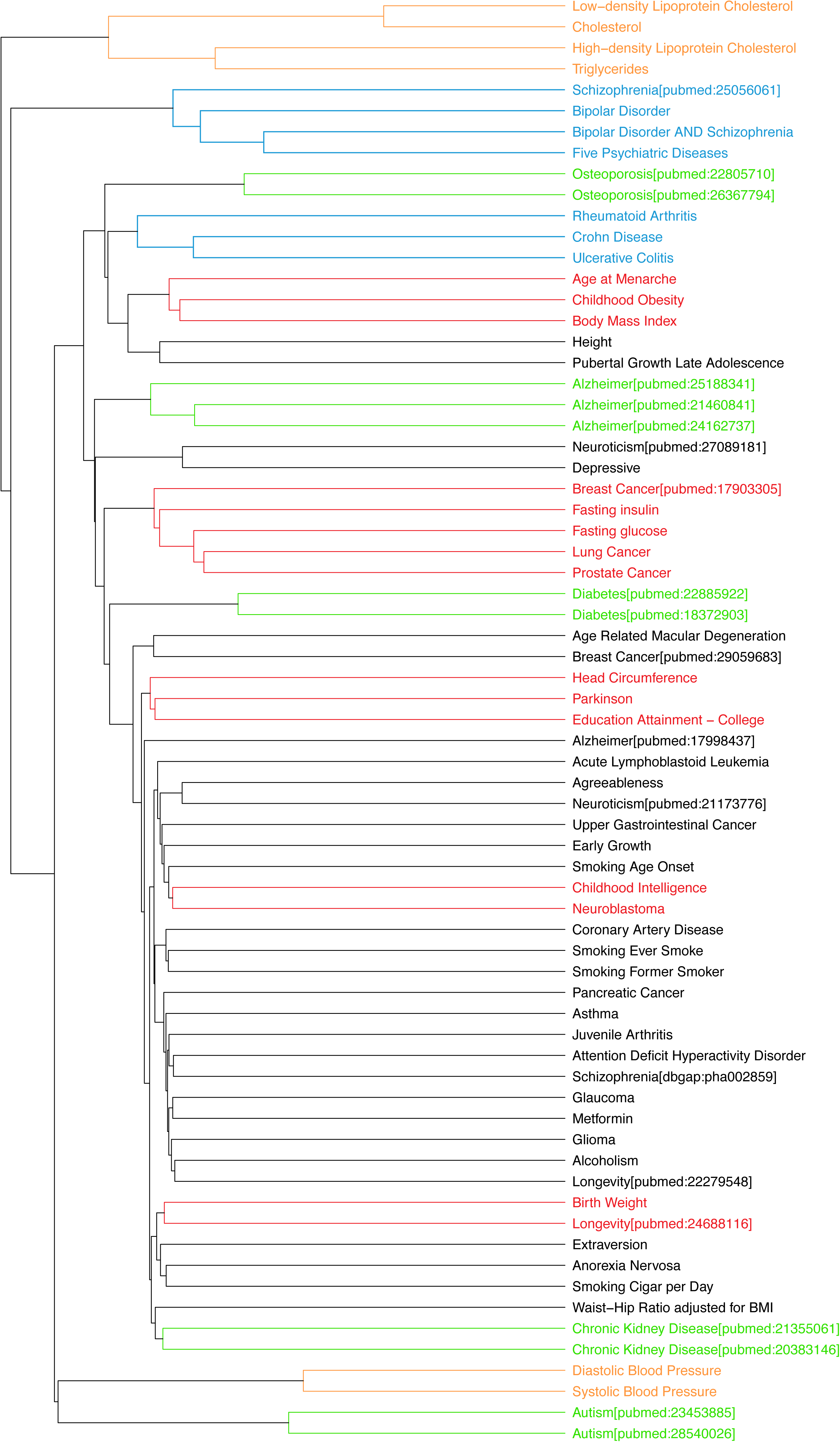
Genetic similarity between phenotypes revealed by the global hierarchical clustering of phenotypes based on their gene association profiles. Highlighted branches discussed in the text: 1) GWASs of the same phenotypes (green); 2) phenotypes that are closely related and analyzed in the same GWAS (orange); 3) phenotypes that are known to share genetic etiology (blue); and 4) phenotypes with no obvious relations but co-occurences were observed based on previous epidemiological studies (red).

The expected phenotype sub-clusters are formed by 1) GWASs of the same phenotype (e.g., osteoporosis, Alzheimer, diabetes, chronic kidney disease, and autism), indicating that at the global level the association was replicated; 2) phenotypes that are closely related and were studied in the same GWASs (e.g., LDL cholesterol, HDL cholesterol, total cholesterol, and Triglyceride; diastolic and systolic blood pressure); 3) phenotypes that are known to share genetic etiology in previous studies (e.g., bipolar disorder (BP), schizophrenia (SZ), the combination of the BP and SZ, and five psychiatric disorders including BP and SZ [35], and phenotypes with strong evidence supporting their shared mechanisms (e.g., rheumatoid arthritis, Crohn’s disease, and ulcerative colitis, due to their common inflammatory/autoimmune nature). The fact that these different groups of related phenotypes were clustered together suggests that the gene-phenotype vector captured the major genetic information of the phenotype.

The global similarity analysis based on the gene-phenotype vector revealed a number of sub-trees in which phenotypes with no obvious relations are linked together (Figure 2, red subtrees). Here we discuss a few examples: 1) a subtree that contains age at menarche, childhood obesity, body mass index; 2) a subtree with 5 phenotypes: breast cancer, fasting insulin, fasting glucose, lung cancer, and prostate cancer; 3) head circumference, Parkinson, and education attainment; 4) childhood intelligence with neuroblastoma; 5) longevity with birth weight. Upon literature search, we found that for each of the groups, there were previous epidemiologic studies that found the co-occurrence of the phenotypes: 1) a previous study found that girls experience early menarche are significantly often overweight/obese [36]; 2) hyperinsulinemia with insulin resistance is a significant risk factor for breast cancer independent of adiposity or body fat distribution [37]; 3) higher educational attainment significantly lower the risk for Parkinson’s disease [38]; 4) children with neuroblastoma score a significantly higher IQ than the control [39] 5) mouse experiment showed that low birth weight correlates with increased longevity [40]. These were the observed correlations of the phenotypes, and the fact that they are clustered by the gene-phenotype association matrix indicates that they are linked by the associations with a set of genes that should shed light on the common underlying genetic mechanisms.

### Delineating common genetic mechanisms underlying seemingly unrelated phenotypes

To identify potential common mechanisms that underlie two seemingly unrelated phenotypes, we search for genes that drive the similarity between the two gene-phenotype vectors. One straightforward approach is to analyze the most significantly associated genes with each of the phenotypes and test for significant overlap. This approach yielded insight for some of the phenotype subtrees. One example is the age-at-menarche/body-mass-index/childhood-obesity subtree. We found overlapping genes in each of the three pairwise comparisons (Figure S3). Multiple genes are associated with age-at-menarche and body-mass-index with a theme on cell growth and proliferation. For example, SMAD3 is significantly associated with both age-at-menarche and body-mass-index and is a pivotal intracellular nuclear effector of the TGF-beta (transforming growth factor beta) signaling which play a critical role in regulating cell grow, differentiation and development[41]. MAP2K5 (Mitogen-Activated Protein Kinase Kinase 5) is also significantly associated with both the phenotypes and the signaling cascade mediated by this kinase is involved in growth factor stimulated cell growth and muscle differentiation [42]. Genes simultaneously associated with both childhood-obesity and body-mass-index include TFAP2B and CENPO. TFAP2B is a transcription factor of AP-2 family which stimulate cell proliferation and suppress terminal differentiation of specific cell types during embryonic development [43]. CENPO encodes a component of the interphase centromere complex which is required for bipolar spindle assembly, chromosome segregation and checkpoint signaling during mitosis thus is critical to cell division[44]. Thus examination of the shared significant gene associations suggests a common theme on cell growth and development underlying all three phenotypes.

While comparing the top associated genes between phenotypes yields clues about the common mechanisms for some of the phenotype subtrees, its power of detection is limited. For many phenotype pairs that are similar based on their gene association p-value vector and thus clustered together, the top associated genes for the two phenotypes do not overlap (an example is the subtree with 5 phenotypes including fasting-insulin and breast-cancer). This suggests that the similarity is driven by more diffused signal distributed to a set of genes with weak associations. To identify such set of genes, we employ an approach previously developed to identify a subset of functionally related genes that drive the similarity between two gene expression profiles[45]. This approach assesses the contribution of a subset of genes defined by a GO category (or a pathway) to the Pearson correlation and assigns a z-score to the subset (see Method). This allows the ranking of the significance of different subsets of genes based on their z-score; we call it Partial-Pearson-Zscore.

Using the Partial-Pearson-Zscore, we analyze the pairs of phenotypes in a subtree and found many significant GO categories that contribute to the similarity, suggestive of common underlying mechanisms for these phenotype pairs. As expected, for the subtree with rheumatoid-arthritis/Crohn’s-disease/ulcerative-colitis, we found that the top GO categories that contribute to the similarity between RA and CD are “antigen processing and presentation of peptide or polysaccharide antigen via. MHC class II” (ranked 1^st^, z-score = 18.6) and similar GO categories (“MHC class II protein complex” ranked 2^nd^, and “peptide antigen binding” ranked 3^rd^), and pyroptosis (an inflammatory form of cell death) (ranked 6^th^, z-score= 14.8). Similarly, GO categories contributing to the similarity between RA and UC include pyroptosis (ranked ^1st^ z-score = 20.7). These results suggest that at least part of the common underlying mechanisms for these diseases are immune in nature and involve inflammation induced cell death.

For the subtrees with phenotypes not obviously related, Partial-Pearson-Zscore analysis also revealed significant GO categories suggestive of the genetic mechanisms that connect these phenotypes together. An example is the subtree with 5 phenotypes including fast insulin and breast cancer (Figure 2). Significant GO categories were found for all 10 pair-wise comparisons (Figure 3A). For example, we found that the “cgmap catabolic process” is the most significant GO category for the fast-insulin/breast-cancer pair. This category includes several nucleotides phosphodiesterase, such as PDE4A and PDE2A. These are dual specificity cAMP and cGMP phosphodiesterase, which degrade cAMP and cGMP, and thus suppress cAMP/cGMP signaling. Inhibitors of these enzymes have been used as anti-cancer drugs[46], as the activation of cyclic nucleotide signaling is sufficient to inhibit proliferation and activate apoptosis in many type of cancer cells[46, 47]. cAMP signaling pathway is also strongly connected to insulin secretion. Among the various intracellular signals involved, cAMP is particularly important for amplifying insulin secretion[48]. Thus altered cAMP/cGMP signaling seems to connect with both breast cancer and fast-insulin.

**Figure 3.**
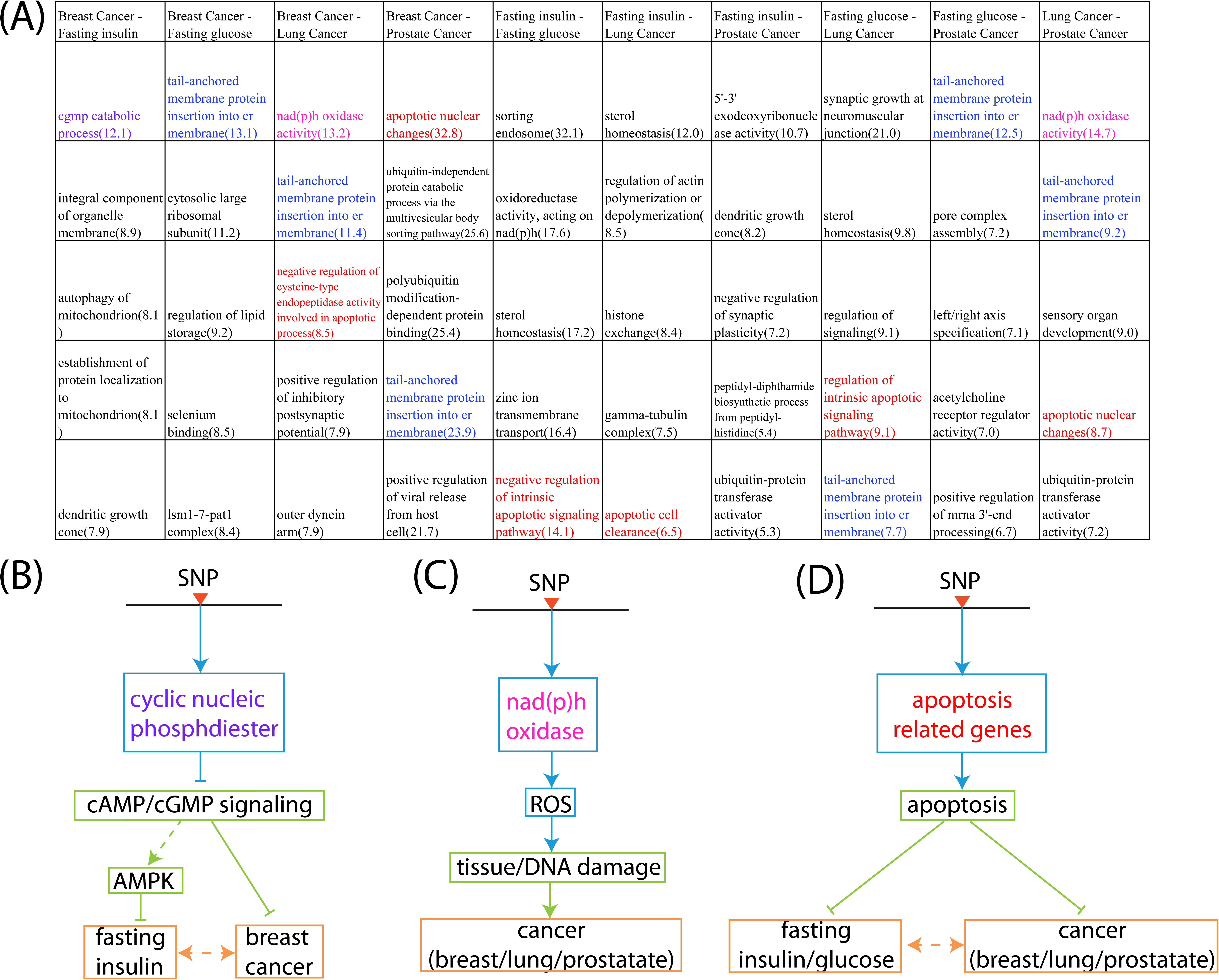
Summary of the top GO categories contributing to the overall similarity between phenotypes for the subtree including five phenotypes (fasting glucose, fasting insulin, breast cancer, lung cancer, prostate cancer). (A): Top 5 non-redundant GO categories with highest Partial-Pearson-Zscore for each pair of the phenotypes. Recurrent GO terms across different pairwise comparisons were highlight with different colors. Also highlighted is the “cgmp catabolic process” that support the relation between breast cancer and fasting insulin. (B) to (D): proposed models with causal relations to explain the observed co-associations between the expression of the genes in the GO term and multiple phenotypes. Alternative models were considered in the discussion section.

Among the most significant GO terms from the pair-wise comparisons, “NAD(P)H oxidase activity” occurred multiple times and connects all three cancer types (breast cancer, prostate cancer, lung cancer). NAD(P)H oxidases are known to express at high level and produce ROS in cancer cells. Over-expression of these enzymes (possibly induced by inflammatory signal) has been linked to tissue injury and DNA damage from ROS that accompany pre-malignant conditions, contributing to the initiation and progression of a wide range of solid and hematopoietic malignances[49].

One of the most frequently appeared GO terms is related to apoptosis (Figure 3A, colored red). This theme occurred in 6 out of 10 phenotype pairs and connects all 5 phenotypes. Defect in the normal programmed cell death mechanisms play a major role in the pathogenesis of tumor [50]. Apoptosis is also known to play an important role in the pathophysiology of both type I and type II diabetes and excess apoptosis is the underlying cause for beta-cell loss [51]. Thus altered apoptosis might be the common mechanism connecting the three cancer phenotypes and changed fasting-insulin/fasting-glucose level in normal people.

Another GO term that connect all the phenotypes is “tail-anchored membrane protein insertion into ER”. This GO term includes several genes (such as SGTA and BAG6) involved in the post-translational pathway through which tail-anchored membrane proteins are recognized and targeted to ER membrane[52]. This pathway is important in many cellular processes including the regulation of cell cycle. SGTA over-expression was involved in the pathogenesis of breast cancer [53] and BAG6 is involved in the control of apoptosis and is required for the acetylation of P53 upon DNA damage, linking them to cancer risk [44]. ER membrane targeting is also important for insulin secretion, potentially connecting these genes to the fasting insulin/glucose phenotype.

We propose models with causal relations to explain the observed co-associations between the expression of a set of functionally related genes and multiple phenotypes (Figure 3B to D). These models are not unique, as there exists alternative and possibly less parsimonious models that can also explain the observed co-associations (see Discussion).

### Two way bi-clustering detects subsets of genes co-associated with multiple phenotypes

The Partial-Pearson-Zscore approach can detect the co-association of a subset of gene defined by a GO term or pathway with a pair of phenotypes, and examination of all the pairwise comparisons can reveal common GO terms/pathways associated with multiple phenotypes clustered in a sub-tree. With the complex relations between physiological processes and phenotypes (Figure 1), it is possible that multiple genes not necessarily belonging to a predefined functional category can co-associate with multiple phenotypes not necessarily in the same sub-tree.

To detect the co-association of a subset of genes with multiple phenotypes, we applied a two-way bi-clustering algorithm – the Iterative Signature Algorithm (ISA) [54] to the global gene-phenotype association matrix. This algorithm was developed previously to analyze gene expression data in order to detect a subset of genes with similar expression patterns across a subset of experimental conditions. The ISA algorithm starts with a seed of a sub-matrix and iterates through phenotypes and genes to recruit/drop members till the set of phenotypes and genes converge to a stable submatrix. Statistical significance of detected phenotype-gene sub-clusters are calculated by random permutation of the input p-value matrix within each phenotype (see Method).

Starting from a filtered global gene-phenotype association matrix to improve the signal and reduce the search space for the ISA algorithm (see Figure 4A and Method), we have identified a number of significant sub-clusters that connect multiple genes with multiple phenotypes. In general, these sub-clusters involve phenotypes that are expected to be together as well as phenotypes that are seemingly unrelated, and genes with a range of different functional categories, suggesting a complex web of connections between physiological processes and phenotypes (Figure 1). Here we discuss the most significant sub-cluster and provide a list of sub-clusters discovered by the bi-clustering algorithm in Figure S4.

**Figure 4.**
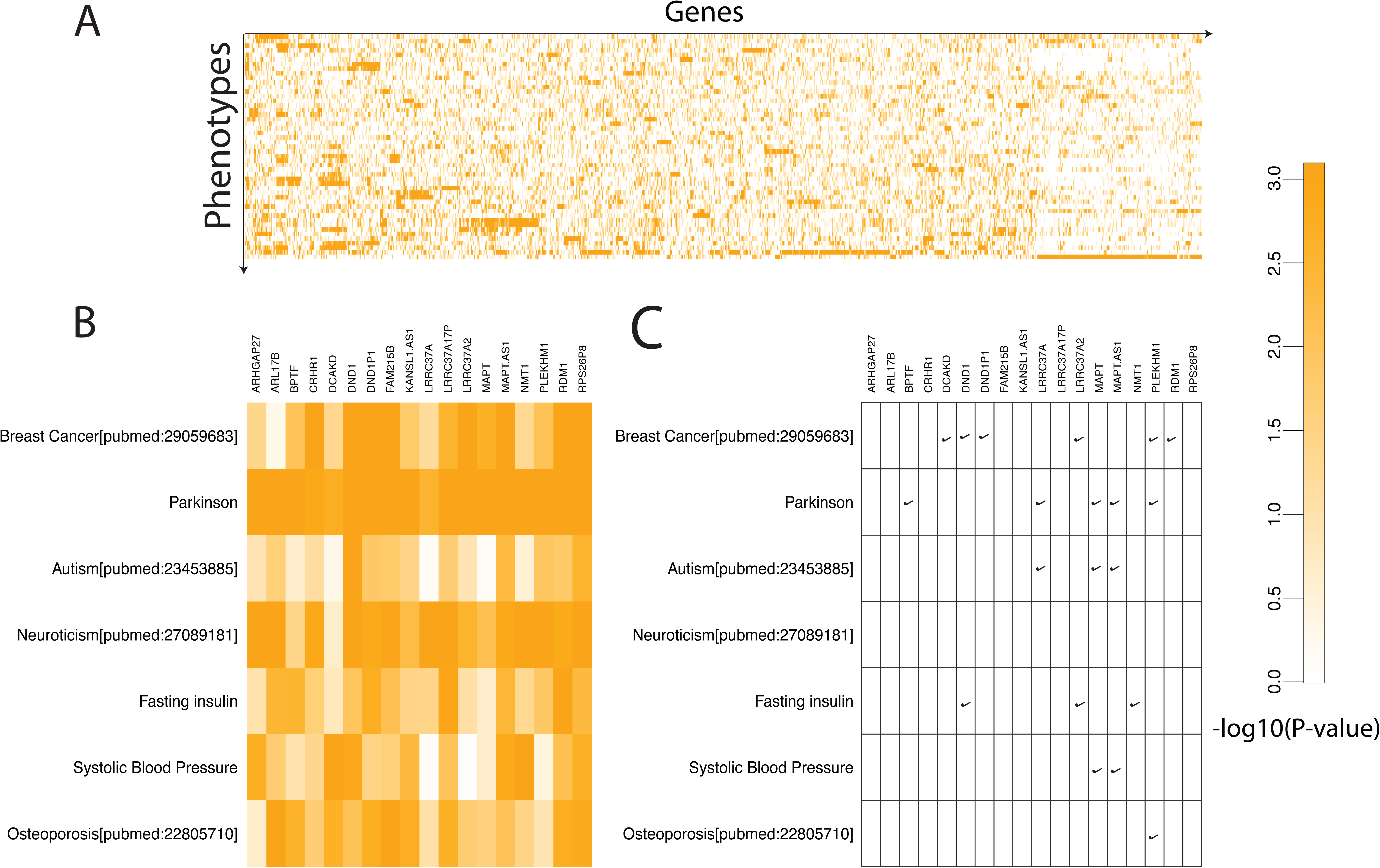
Subclustering analysis revealed a set of genes co-associated with multiple phenotypes. (A) the global gene-phenotype association matrix, filtered to enrich the signal and reduce the dimension. The rows and the columns of the filter matrix were reordered by the hierarchical clustering algorithm. Certain subclusters are already visible. (B) One of the sub-clusters identified by the ISA algorithm. This subcluster consists of 18 genes and 7 phenotypes including a few neuro-psychiatric diseases, osteoporosis, breast cancer and physiological traits such as insulin level and blood pressure. (C) gene-phenotype associations in the subcluster supported by literature (check marks).

The most significant subcluster contains 7 diverse phenotypes and 18 genes with a broad range of biological functions (p-value = 1E-6 Figure 4B). According to gene annotations [44], this set of genes seems to be enriched for regulatory function: ARHGAP27 is a Rho GTPase activating protein, CRHR1 encodes a corticotropin-releasing-hormone-receptor, BPTF is a transcription factor highly expressed in the brain of Alzheimer’s patients, DND1 is an inhibitor of miRNA mediated repression, FAM215B is a non-coding RNA, KANSL1 is a component of histone acetylation complexes, and PLEKHM1 is an adaptor protein that regulates Rab7 dependent fusion events in the endolysosomal system. Some of the gene-phenotype associations predicted by this subcluster are supported by literature (Figure 4C and table S3), and several genes associate with multiple phenotypes across the subcluster. For examples, MAPT (microtubule associated protein tau, known to be associated with multiple neurodegenerative and neuropsychiatric diseases [55]) is associated with Parkinson[56], Autism[57], and Systolic blood pressure [58]. Consistently we found MAPT antisense transcript with similar association patterns, as the expression of MAPT antisense is negatively correlated with MAPT expression [57, 59–62]. PLEKHM1 is known to be associated with breast cancer, Parkinson, and osteoporosis, and mutations in this gene are associated with autosomal recessive osteopetrosis type 6 [63–66]. These examples suggest that other gene-phenotype associations in this submatrix may also be true and biological meaningful associations.

Among the seven phenotypes in this subcluster, several were known to be correlated based on previous epidemiological studies. For example, in addition to the connection between fasting insulin and breast cancer discussed above, insulin was linked to several phenotypes in this cluster: 1) there is a high prevalence of insulin resistance in non-diabetic Parkinson’s disease patients [67]; 2) it was suggested that abnormal insulin signaling underlies increased autism risk based on the evidence that mutations that hyperactivate PI3K pathway cause autism [68]; 3) insulin resistance co-occurs with hypertension[69]; 4) insulin may work to stimulate osteoblast differentiation, and diabetes have a higher risk of bone fracture[70]. These evidences suggest that insulin signaling may be one of the physiological processes that connect these phenotypes together.

Functional analysis of genes in the subcluster added further support that regulation of insulin signaling is a common mechanism, with three of the genes in the cluster having strong connection with insulin. The first gene CRHR1 encodes a corticotropin-releasing-hormone-receptor. Administration of corticotropin, the downstream functional hormone of CRHR1, results in a 5-to 10-fold rise in plasma insulin levels in mice [71]. In ER positive breast cancer which constitutes about 80% of all breast cancers, estrogen alters the splicing of CRHR1 and disrupts CRH-mediated signaling, which contributes to the cancer growth driven by estrogen[72]. Thus CRHR1 may connect insulin and breast cancer through the downstream insulin signaling. The second gene is NMT1(N-myristoyltransferase). Its association with insulin is supported by mice experiment that liver particulate N-myristoyltransferase activity appears to be inversely proportional to the level of plasma insulin [73]. The third gene is PLEKHM1 which is an adaptor protein that regulates Rab7 dependent fusion events in the endolysosomal system[44]. PLEKHM1 participates in the degradation of insulin granule by the fusion of secretory granule with lysosomal, which is highly up-regulated upon starvation [74]. Thus loss of function mutations in this gene may lead to high insulin level that drives osteopetrosis.

## Discussion

We have developed a computational framework to study the genetic similarity between different phenotypes and to delineate the potential mechanisms underlying the similarity by integrating GWASs of multiple phenotypes with association studies of IMT. We demonstrated the approach by a global analysis of 445 GWASs covering 92 complex human phenotypes. Our analysis revealed expected as well as unexpected relationships between phenotypes, and suggested specific mechanisms that connect these phenotypes together. Our analysis also revealed many specific gene-phenotype and metabolite-phenotype associations that should motivate more detailed mechanistic studies in the future.

Comparing with genetic similarity analysis using GWAS data at the SNP level, our analysis framework has several advantages. First, comparing the genetic profiles at the gene level can yield stronger signal, as multiple SNPs can converge on the same gene. Even if the GWAS peaks of two different phenotypes do not align by proximity, they can be “aligned” through a gene, i.e., they may influence the expression of the same gene. We provided one example in which the gene RPL23 is co-associated with both schizophrenia and height (Figure S2); the correlation of the two phenotypes has been observed in epidemiological studies [13]. The co-association is supported by two GWAS SNPs associated with the two phenotypes respectively. These two SNPs do not align by proximity; they align to two different eSNPs for the same gene.

Secondly, genetic similarity analysis at the gene level readily suggests potential mechanisms connecting different phenotypes. We have presented two different approaches to delineate genes/pathways that contribute to the genetic similarity: the Partial-Pearson-Zscore analysis to identify a group of functionally related genes that contribute to the similarity between a pair of phenotypes, and a sub-clustering analysis of the global gene-phenotype association matrix that identifies a set of genes co-associated with multiple phenotypes. Analyses using these approaches led to specific hypotheses regarding the mechanisms that connect expected as well as unexpected relationships between phenotypes, and we expect that the power of detection will grow as more GWAS and IMT data accumulate.

It is important to note that the data we obtained using this analysis framework are co-associations between genes and phenotypes supported by multiple GWAS and eQTL SNPs, not causal relations. In suggesting mechanistic models, we proposed specific causal relationships that are consistent with the data (Figure 3). However, there are often alternative (and perhaps less parsimonious) models that are consistent with the observed data. For example, we proposed that SNPs influence genes involved in apoptosis which can independently influence fasting insulin/glucose levels and cancer risks (Figure 3D). However, there exists alternative models that are also consistent with the data. For example, SNPs can influence fasting insulin level which in turn influences apoptosis associated genes’ expression which in turn influences cancer risk. A variation of this model is that insulin can influence apoptosis associated genes’ expression and cancer risk independently (Figure S5). These alternative models are also consistent with the co-associations between SNPs and all the phenotypes (including the gene expression phenotype), but less parsimonious in that there is a missing molecular trait in between SNPs and fasting insulin/glucose. However, the true mechanistic model is not necessarily the most parsimonious. Clearly more experiments/analyses are needed in order to discriminate between different models.

## Method

### Sherlock-II algorithm

Sherlock-II detects IMT-phenotype association by comparing the GWAS SNPs with the QTL SNPs of the IMT (eQTL or metabolite-QTL), with the assumption that if an IMT is causal to the phenotype, SNPs that influence the IMT should also influence the phenotype, thus the QTL profile of the IMT should have significant overlap with the GWAS profile of the phenotype. To test the significance of the overlap, Sherlock-II first selects the QTL SNPs of an IMT that pass a threshold, aligns these SNPs to the corresponding SNPs in the GWAS, and test whether the aligned GWAS SNPs have p-values significantly better than those picked at random.

When testing the significance of a selected set of loci in a given GWAS, Sherlock-II uses a simple scheme to compute a score for the n p-values of the GWAS SNPs that best tag these loci. It then uses a convolution-based approach to construct an empirical null distribution for scores involving n p-values drawn randomly from the full set of GWAS p-values. In certain instances, particularly with eQTL for many genes, this test will be performed repeatedly for all intermediate molecular traits in a study. If dependence between the sets of IMTs is detected — for example, due to pleiotropic loci that regulate many genes — a provision of adjusting the null distribution is included. Through extensive testing and simulation, we show that our empirical strategy produces p-values that are highly resistant to inflation. In this section, we describe the individual components of the Sherlock-II method in detail in the order in which they occur.

#### Identifying Tag SNPs for Independent Blocks

Sherlock-II statistics is based on the assumption that the collection of SNPs tested are tagging independently segregating haplotype blocks in both the GWAS and eQTL cohorts. Understanding the true block structure of the cohorts is critical to accurately estimating gene significance. The inadvertent inclusion of dependent blocks can inflate the significance of genes by over-counting true associations; conversely, a conservative approach that enforces wide separation between tagging SNPs may miss many independent loci. The preferred approach to identifying independent blocks requires genotypes for both the GWAS and functional data to define block boundaries based on regions of rapid decay in linkage-disequilibrium (LD). However, since the difficulty of obtaining genotypes across a large number of GWAS studies makes this impractical, we constructed a database of LD using Caucasian cohorts from the 1000 Genomes Project. We use PLINK (v1.07) to compute r^2^linkage between common SNPs for 379 individuals in the CEPH, CEU, TSI, GBR, FIN, and IBS cohorts of 1000 Genomes release v3.20101123. This permits the alignment of associated loci in the functional data to corresponding SNPs in GWAS data, and it enables the identification of independently-segregating tag SNPs in the combined data for use in the statistical test.

We first match the GWAS SNPs to all the loci present in a given functional data set that pass a specific threshold. For the eQTL data used here, the nominal threshold is typically eSNP p-values < 10 ^-5^. If matching SNPs are not present in both data sets, we use the closest GWAS SNPs in r^2^ LD > 0.85, if available. We then use an agglomerative clustering approach to identify non-overlapping blocks of SNPs in r^2^ LD < 0.2. From each block, the functional SNP with the strongest association is selected as a tag, and its corresponding GWAS SNP is included in the set subjected to statistical test. Since discrepancies between the various cohorts — GWAS, functional, and linkage — are inevitable, a minimum 100 kb distance between tag SNPs is enforced. In addition, we exclude SNPs from the human leukocyte antigen (HLA) region between 6p22.1 to 6p21.3 due to its complex linkage structure.

#### Core Statistical Method

Once a set of SNPs related to a specific molecular trait is identified, we compute a score s by simply combining their GWAS log p-values:

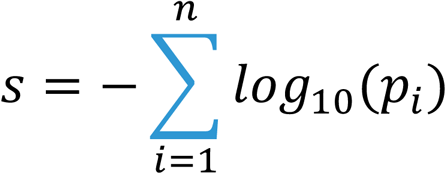

This converts our collection of SNPs into a scalar quantity that, when referenced against the appropriate null distribution, indicates the statistical significance of selecting this subset from the pool of all independent GWAS loci. Our approach is analogous to Fisher’s combined probability test using the scoring function 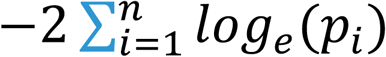 which follows a χ^2^ distribution with 2n degrees of freedom when p-values are drawn from a uniform distribution on (0,1). With our method, instead of assuming a particular form for the distribution of GWAS p-values, we use discrete convolution to compute an empirically-derived distribution of scores when combining n p-values. Since s represents a specific score, we let S represent a random variable of all possible scores. We form the discrete probability distribution function (PDF) of S using bins of width b log units, where the probability of scores in the range 0 ≤ s < b is the first element, b ≤ s < 2b is the second element, and so forth. Thus, f_n_[s] = P(S = s) is the discrete PDF for a score comprising n independent GWAS p-values. For the simplest case of scores involving only a single SNP, the PDF f1[s] is essentially a normalized histogram of p-values for all independent (unlinked) SNPs in the GWAS that aligned to a eQTL SNP across all genes:

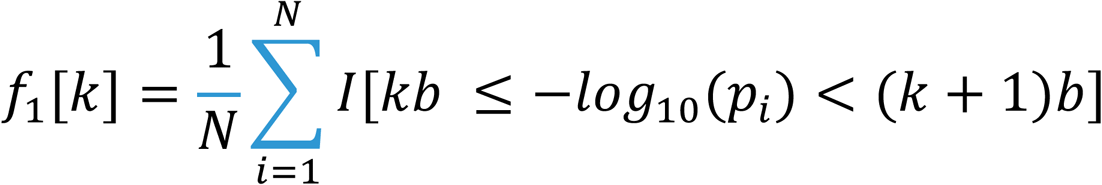

where I is the indicator function, N is the total number of SNPs, b is the bin width, and k = s/b is the array index. For computational efficiency, the minimum GWAS p-value is typically truncated at 10 ^-9^, well below genome-wide significance, yielding 900 bins when spacing b = 0.01 is used. This single-locus PDF forms the basis from which null distributions for any arbitrary number of functionally-related loci are formed. Since the sum of two independent random variables has a PDF equal to the convolution of their individual probability distributions, scores involving two GWAS p-values have PDF f_2_ = f_1_ * f_1_. In the general case for any value of n > 1, the PDF f_n_[s] is formed from n-1 recursive convolutions of f_1_[s]:

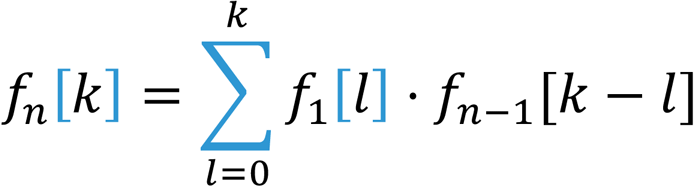

where again k = s/b is the array index. For the special case of a truly uniform p-value distribution, we have an exact continuous solution to the recursive convolution for n p-values (which can be obtained from the chi-square distribution)

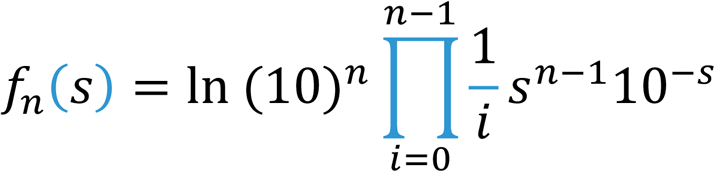

To motivate the use of our empirical approach, we compare this theoretical result to our empirical method for GWAS of differing power. For low-powered association studies with p-value distributions close to uniform, the tail of the empirical distribution is similar to the tail of the theoretical distribution, leading to roughly identical estimates of significance for a given score. For well-powered meta analyses that appear to contain inflation from various sources, significant differences in the distribution tails can lead to an appreciable overestimation of significance for high scores when using the theoretical distribution. With the empirical distribution derived from the real GWAS data, Sherlock-II is insensitive to the input GWAS inflation.

#### Correcting for Pleiotropic and Sampling Effect

Another source of inflation stems from a lack of true independence between molecular traits across a given functional dataset. For example, pleiotropic loci may appear to regulate hundreds of genes in the eQTL data for a given tissue. In our simulations, these loci may inflate the output test statistic due to chance alignment with significant GWAS SNPs; in our tests, this occurs at an appreciable level in perhaps five percent of the cases. Since our method permits the use of a unique distribution f_1_ for each p-value added to the score, it enables a simple scheme for reducing inflation by adjusting each distribution based on the number of molecular traits affected by the locus. For the eQTL example, this involves conditioning the GWAS distribution based on the number of genes influenced by a locus across the entire eQTL data set. When chance alignment of pleiotropic loci and significant GWAS SNPs occurs, the overrepresentation of small p-values is reflected in the null distribution for affected genes. Thus, in practice, the actual distribution incorporated into a Sherlock-II score with eQTL data is conditioned on the number of genes that are regulated by the same locus:

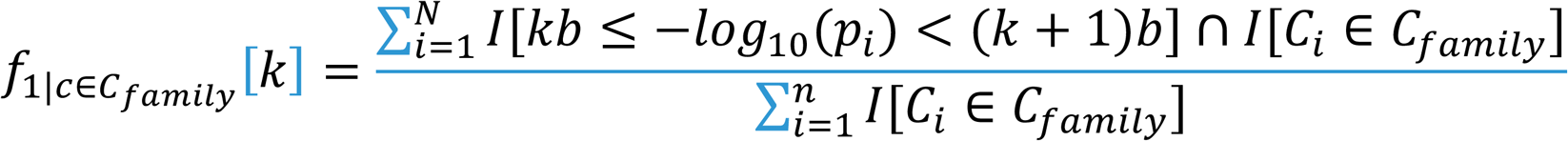

where I is the indicator function, N is the total number of SNPs, k is the index, b is the bin width, and the ith SNP is assigned gene count C_i_ while the locus in question has count c. *C*_*family*_ is several non-overlapping sets, each including a range of gene count c. For example *C*_*family6*_ includes all SNPs that regulate 6 to 10 genes. Now the distribution contains only the p-values of SNPs that belong to the same count family. With this change, instead of computing the significance of a set of SNPs drawn from GWAS, our method computes the significance of a set of SNPs drawn given their individual pleiotropies count family. In our simulations, this results in a minor change in the significance and rank order of output genes in most cases but prevents inflation whenever chance alignment of significant GWAS SNPs and pleiotropic loci occurs (Figure S6). Specifically, we penalize SNPs regulating the expression of a large number of genes.

### Calculate Genetic similarity using gene association profiles for the phenotypes

We first filtered the gene-phenotype association matrix to eliminate redundancies. For each phenotype with multiple GWAS dataset from overlapping cohorts, we chose a representative one with the strongest signal, defined as the one with the largest number of genes with p-value of association smaller than 0.001. Note that the representative GWAS doesn’t have to be the one with the biggest sample size because some signal is only statistically significant in a sub-population.

For genetic similarity analysis, we transformed the p-values in the gene-phenotype association matrix to −log10(p-value), so that genes with small p-values can have a stronger weight in the distance calculation.

Hierarchical clustering is applied to the gene-phenotype association matrix to group phenotypes together based on their genetic similarity. We used R and its default distance measure, the Euclidean distance, to calculate distance between phenotype p-value vectors. We then use Ward’s method to cluster the phenotypes [75]

### Partial-Pearson-Zscore analysis to identify sets of functionally related genes that contribute to the genetic similarity

We employed a simple statistical approach to detect gene ontology terms that contribute significantly to the genetic similarity between two different phenotypes. Gene Ontology gene annotations were downloaded from Ensembl. We removed the GO terms with less than 5 genes (not enough statistics) and more than 100 genes (too non-specific) from the analysis, after which 6796 GO terms were used. For a specific pair of phenotypes, we calculate a Partial-Pearson-Zscore for each of the 6796 GO terms using the following formula (previously developed for analyzing similarity between two gene expression profiles [45])

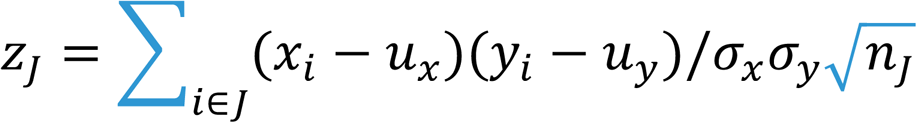

where xi and yi are −log10(p-value) for gene i associated with the two phenotypes, *u and σ* are the mean and standard deviation for x and y respectively, J is the gene set defined by the GO term, and *n*_*J*_ is the number of genes in the GO term. This formula is similar to the one for calculating Pearson correlation (with a normalization 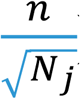), except that the summation is partial –only for the genes in the subset. We argued that the null distribution for *Z*_*j*_ should be standard normal distribution, and verified that this is indeed the case by randomly sample all phenotypes pairs and generate *Z*_*j*_ for all the GO terms (Figure S7).

### Two-way bi-clustering to detect subclusters in which a set of genes associate with multiple phenotypes

In order to enhance signal and reduce the search space, we further filtered gene-phenotype p-value matrix before sub-clustering algorithm is applied. We removed the columns and rows in which none of the gene-phenotype associations passed a Q-value cutoff of 1/3. Thus we only keep the phenotypes and genes with at least one significant association.

We used the ISA algorithm (Iterative Signature Algorithm) [54] to detect sub-clusters. This algorithm was developed to detect a subset of genes with similar expression patterns in a subset of conditions. The ISA algorithm starts with a seed of a sub-matrix and iterates through phenotypes and genes to recruit/drop members till the set of phenotypes and genes converge to a stable submatrix. By default, ISA randomly choose columns/rows as seed to begin iteration with the assumption that a large number of random seeds will cover most starting conditions that converge to stable submatrices. We went further by eliminating the randomness from seeding with an exhaustive set of seeds. More specifically we generated n(n-1)/2 number of seeds for n phenotypes to exhaust all possible combination of starting conditions. Standard post-processing steps with R ISA package [76] are conducted to merge similar sub-clusters.

Statistical significance of the discovered sub-cluster is calculated by permutation test. One permutation is defined by permuting the p-value vector of each phenotype in the gene-phenotype p-value matrix, keeping the p-value distribution within each phenotype invariant. Then ISA algorithm is applied with the same parameters used in the discovery phase to detect sub-clusters from the permutated matrix. A robustness score defined by ISA[54, 77, 78] is calculated for each sub-cluster discovered from the permutated matrix. The highest robustness of these sub-clusters is recorded for each permutation. Sub-cluster p-value is calculated by comparing its robustness score with the distribution of the robustness score generated from all the permutations.

## Figure legends

Figure S1

Four examples of metabolite-phenotype associations identified by applying Sherlock-II to GWAS data and metabolite-QTL data. QQ plot for metabolite p-value distribution (left panel), QQ plot of original GWAS study (inserts in the left panel), and alignment between the metabolite-QTL and GWAS profiles (right panel) are presented. (A) Methionine sulfone is associated with chronic kidney disease. (B) Palmitoyl-dihydrosphingomyelin is associated with Alzheimer’s disease. (C) Gamma-glutamylglutamate is associated with schizophrenia. (D) Homocitrulline is associated with Autism.

Figure S2

(A) Genetic similarity between schizophrenia and height supported by co-association with multiple genes. (B) the gene RPL23 is co-associated with both phenotypes, supported by eSNPs that aligned to non-overlapping GWAS peaks (highlighted with red circles) in two different phenotypes.

Figure S3

Overlapping genes in the three pair-wise comparisons of the phenotypes in the subtree (age-at-menarche/body-mass-index/childhood-obesity) from the hierarchical clustering of the phenotypes.

Figure S4.

Other examples of the sub-clusters discovered by applying the ISA bi-clustering algorithm to the gene-phenotype association matrix.

Figure S5

Alternative models explaining the co-associations among the expression of apoptosis related genes, fasting insulin/glucose, and cancers. (A) the model we proposed in Figure 3. (B)-(C): alternative models that also explain the observed co-associations.

Figure S6

Using p-value distribution conditioned on the pleiotropic counts of the SNPs prevents the inflation of the gene-phenotype association p-values. Top Row: QQ plots from case-control randomization GWASs show no signal. Middle Row: The chance overlap of pleiotropic loci in eQTL data and low p-value SNPs from the randomized GWAS can result in inflation of gene-phenotype association p-values. Bottom Row: the use of PDFs conditioned on SNP pleiotropy corrected the inflation.

Figure S7

Distribution of the Partial-Pearson-Zscore generated by analyzing GO terms for all the phenotype pairs. The distribution has a mean of 0.066 and a standard deviation of 1.06, which are very close to that from the standard normal distribution as expected from a z-score.

