## Supplementary figures and images for "An integrative analysis of GWAS and intermediate molecular trait data reveals common molecular mechanisms supporting genetic similarity between seemingly unrelated complex traits"

### Figure S1

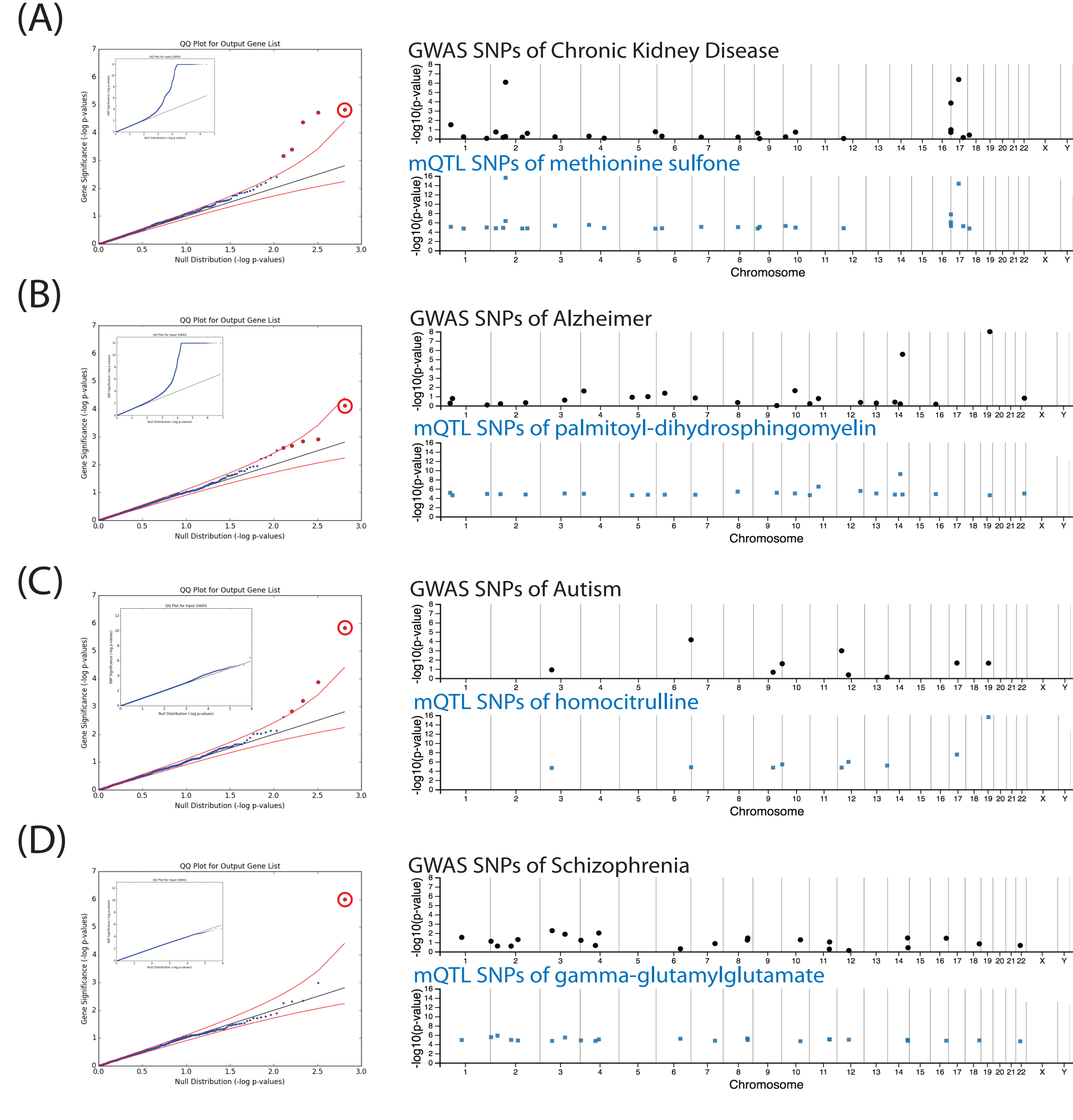

### Figure S2

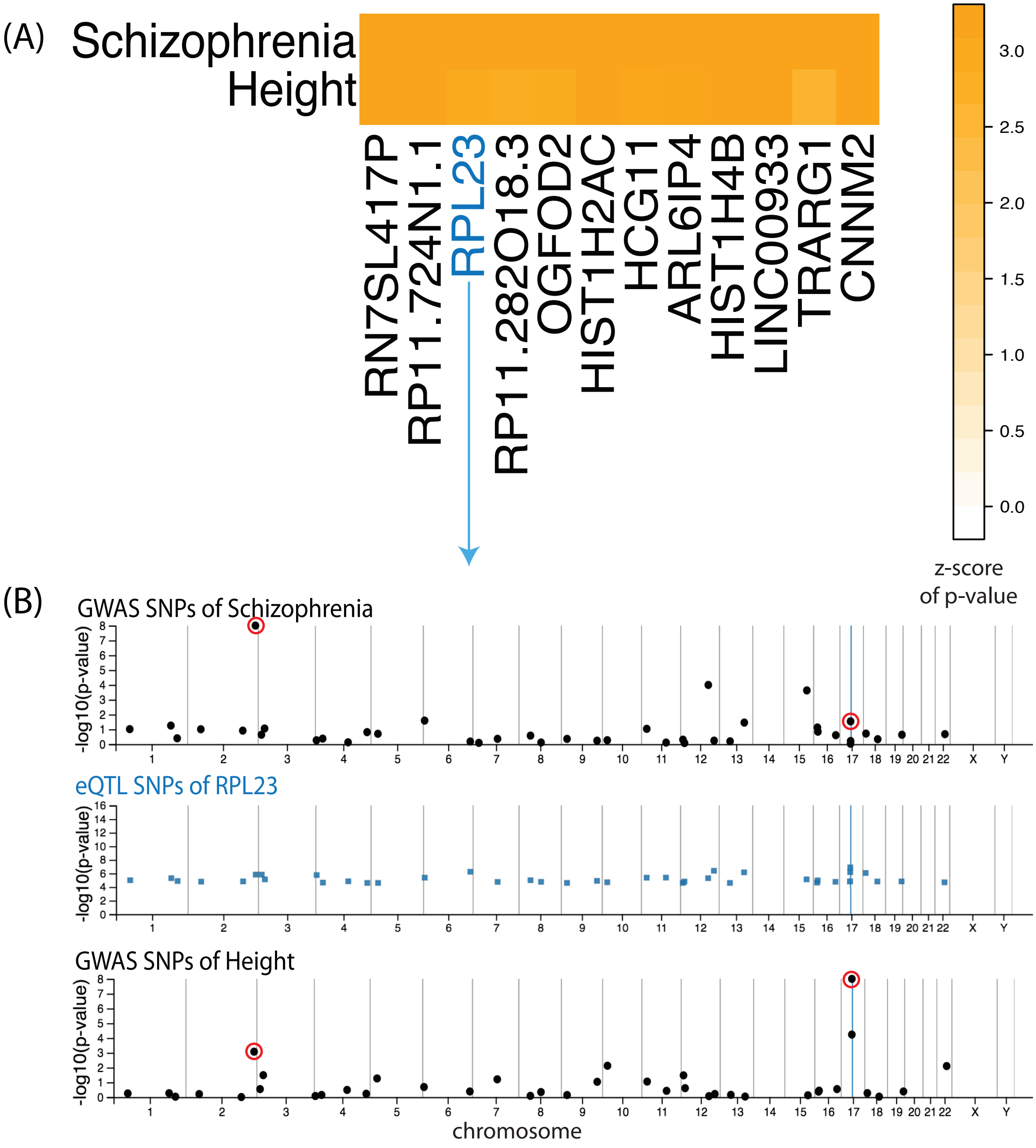

### Figure S3

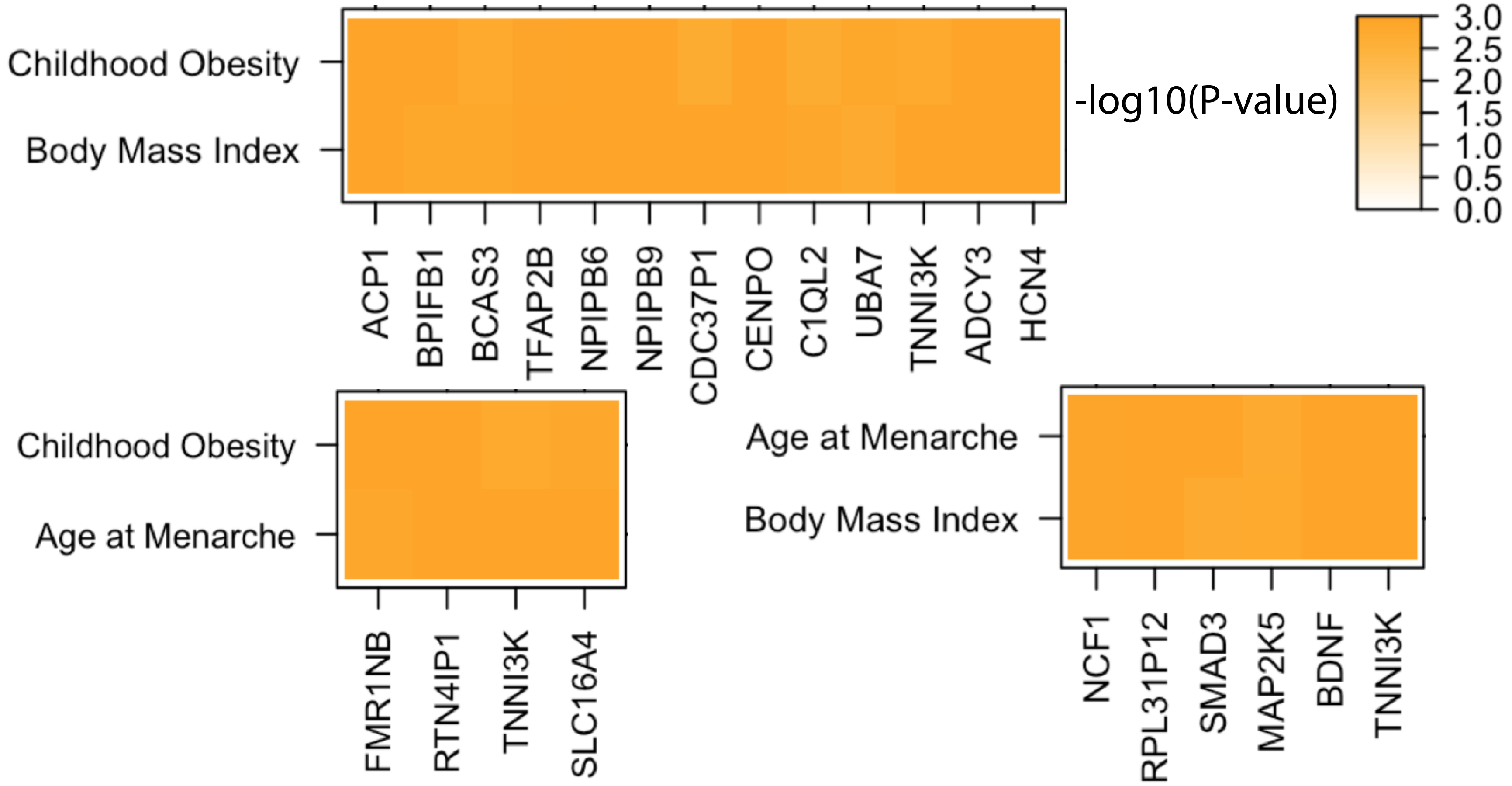

### Figure S4

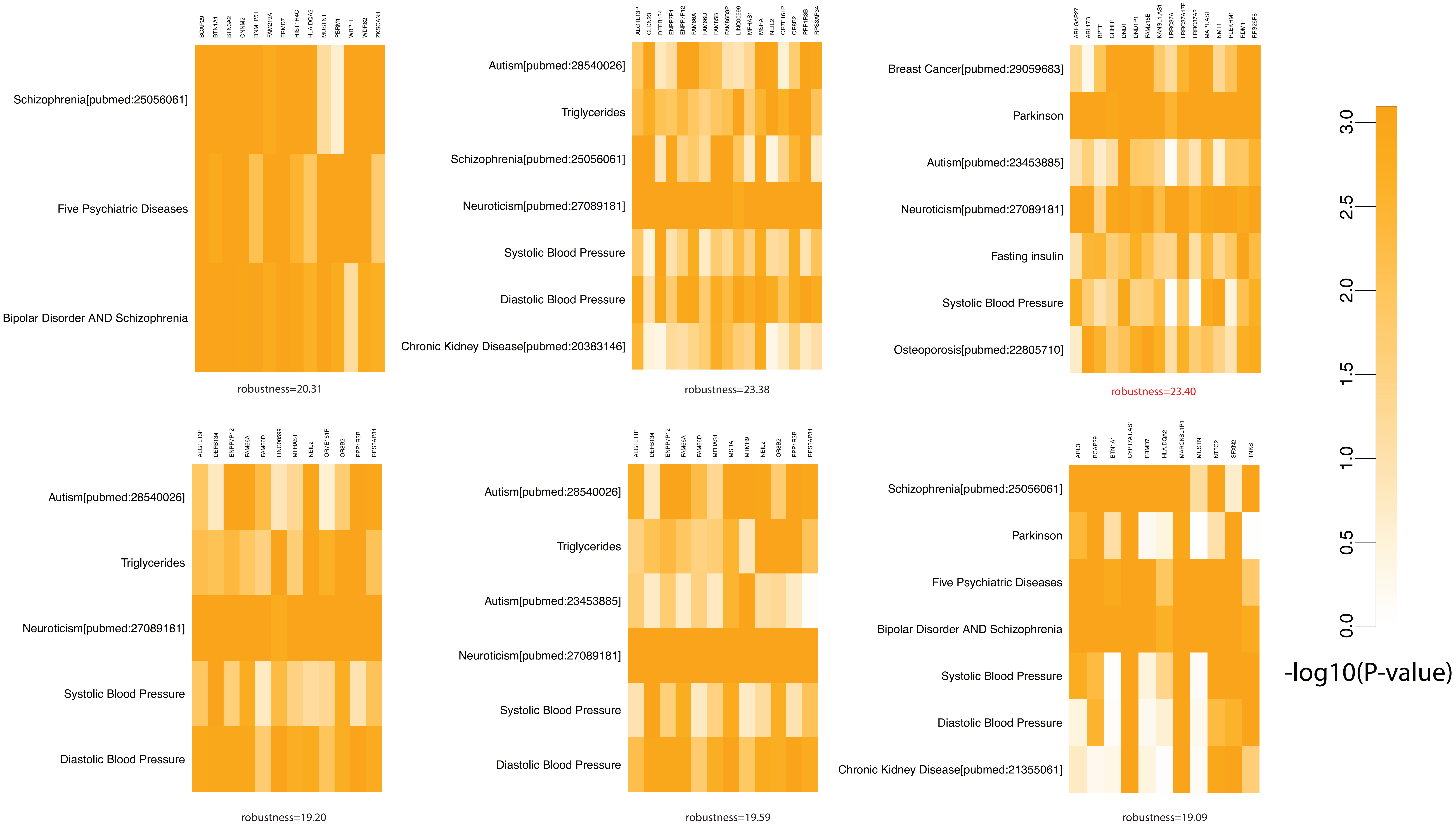

### Figure S5

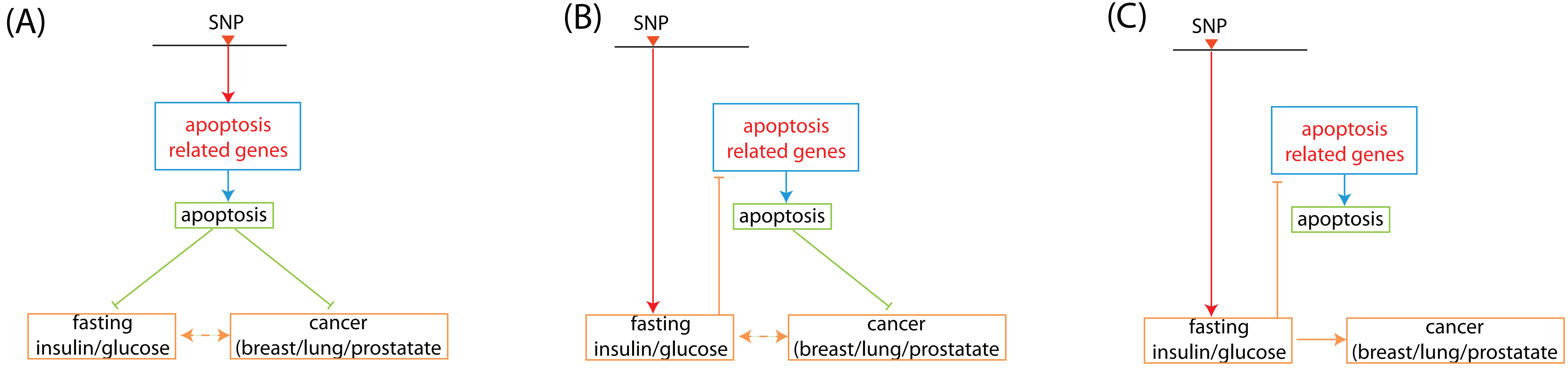

### Figure S6

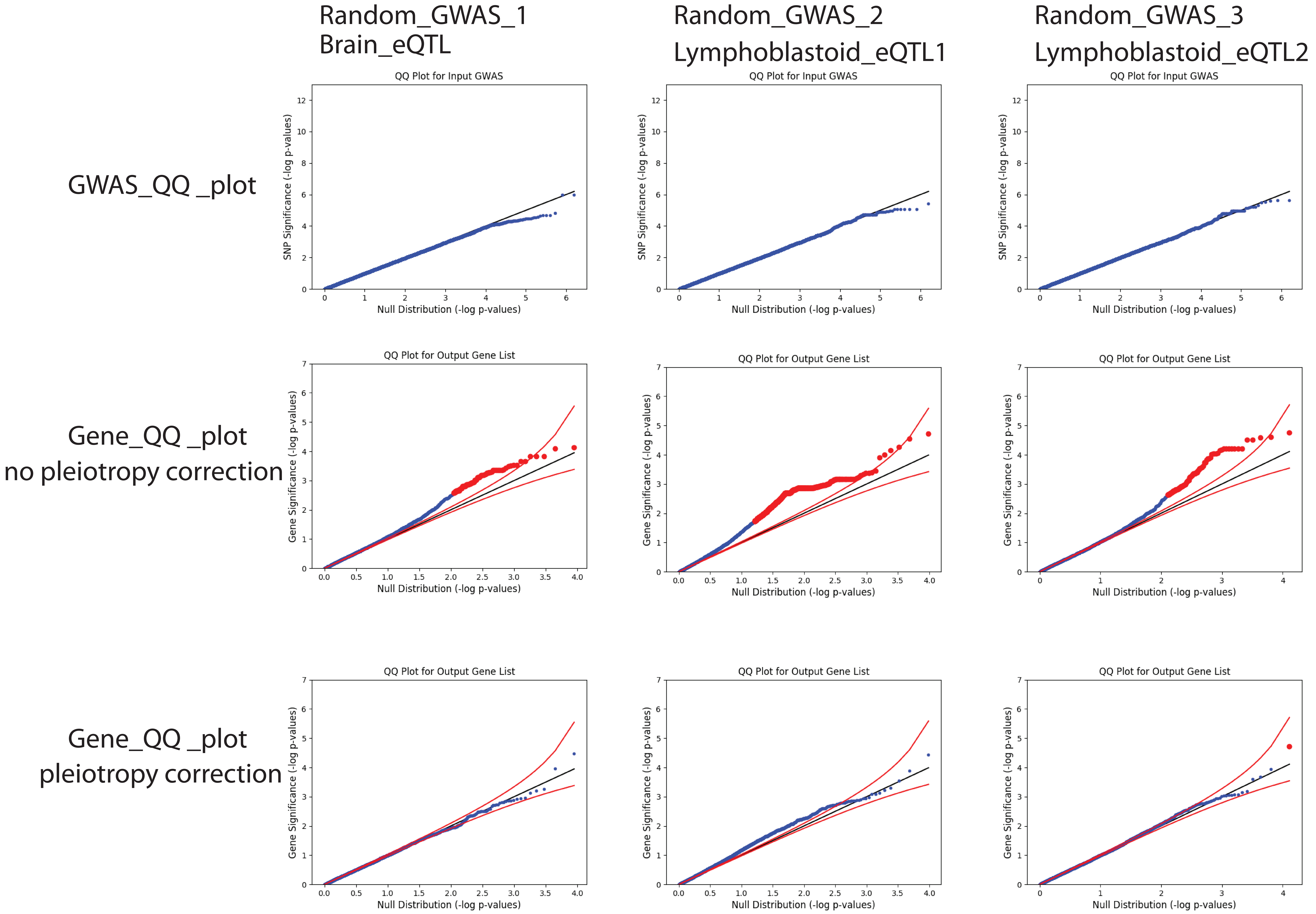

### Figure S7

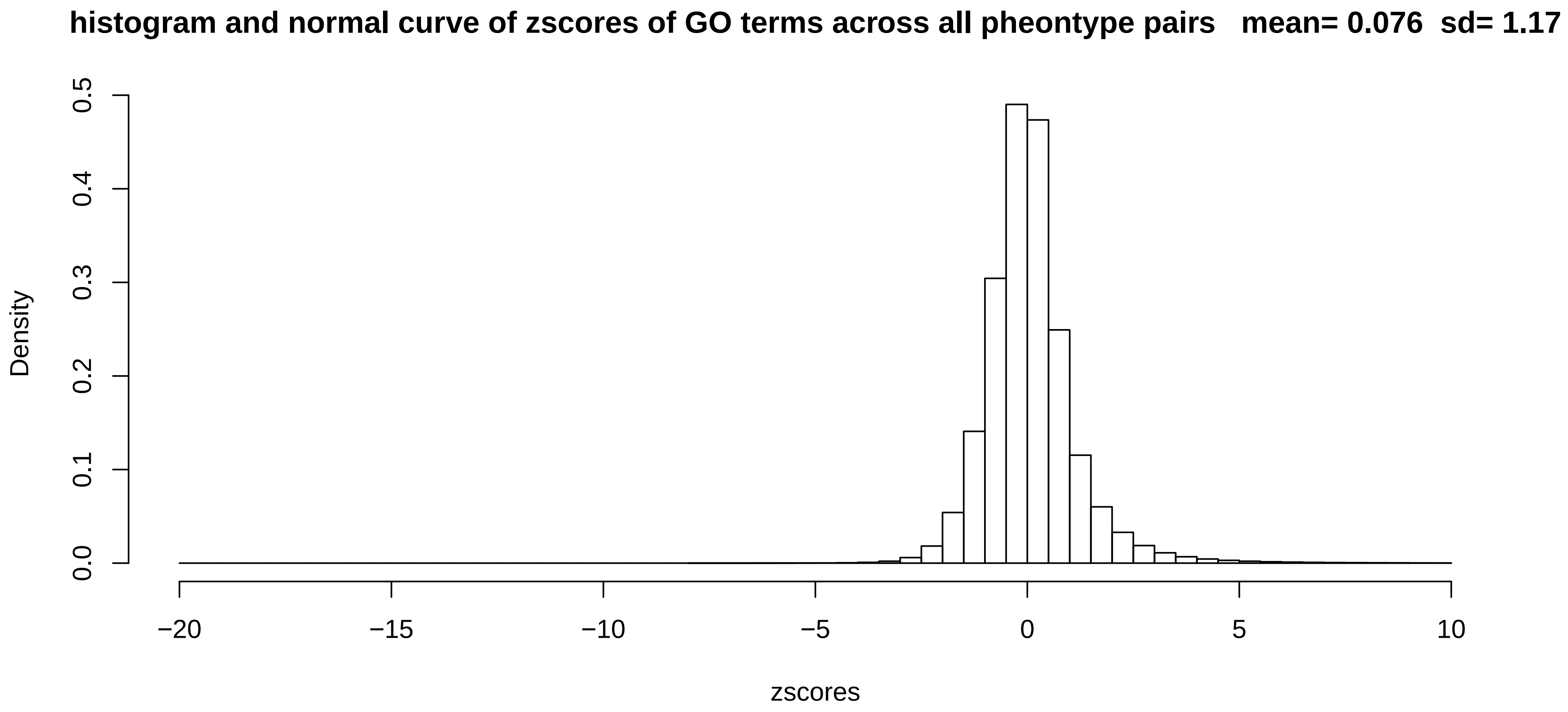
